# A framework for decolonising and diversifying biomedical sciences curricula: rediscovery, representation and readiness

**DOI:** 10.1101/2023.08.15.553224

**Authors:** Tianqi Lu, Zafar I. Bashir, Alessia Dalceggio, Caroline M. McKinnon, Lydia Miles, Amy Mosley, Bronwen R. Burton, Alice Robson

## Abstract

To date, most efforts to decolonise curricula have focussed on the arts and humanities, with many believing that science subjects are objective, unbiased, and unaffected by colonial legacies. However, research is shaped by cultural and historical context, and inequities exist in funding, publishing and acknowledging scientific achievements. Our curricula reflect these inequities, perpetuating bias to future generations of scientists. We examined attitudes and understanding towards decolonising and diversifying the curriculum among students and teaching staff in the biomedical sciences at the University of Bristol, to discover whether our current teaching practice is perceived as inclusive. We used a mixed methods study including surveys of staff (N=71) and students (N=121) and focus groups. Quantitative data showed that staff and students think decolonising the curriculum is important, but this is more important to female respondents (p<0.001). Students are less aware than staff of current efforts to decolonise the curriculum, while students from minority ethnic groups feel less represented by the curriculum than white students. Thematic analysis of qualitative data revealed three themes that are important for a decolonised curriculum: rediscovery, representation and readiness. We propose that this ‘3Rs framework’ could guide future efforts to decolonise and diversify the curriculum in the biomedical sciences and beyond.

## Introduction

Decolonisation of the curriculum represents a conscious shift towards confronting and rectifying the Eurocentric bias in research and curriculum [1]. It challenges the preconceived Eurocentric narrative and other wrongs rooted in systemic societal prejudice. In the aftermath of the “Rhodes Must Fall campaign”, decolonising the curriculum has become a pressing priority, aiming to enable scholars to acknowledge and ameliorate the structural and epistemic legacy of colonialism [2].

Scientific research and education are widely believed to be founded upon objective, evidence-based theories. However, these disciplines originate from the same historical setting that spawned colonialism and its associated racial inequality [3, 4]. European scientific institutions played a key role during colonial times, serving as instruments for colonisation and reaping significant rewards from imperialist exploitation [5, 6]. They are therefore beholden to assist the decolonial endeavour, for example, by retrieving and reconnecting indigenous knowledge that has been erased and discussing problematic practices or figures in the curriculum. This is especially true for the University of Bristol, whose host city was instrumental in Britain’s colonial history regarding the transatlantic slave trade: a recent report commissioned by the University of Bristol discusses this legacy as it works towards positive change [7]. Bristol became a particular focus of attention in 2020, when a statue of the slave trader Edward Colston was torn down during an anti-racism protest in the city [8].

Scholarly work in decolonisation is not new, with advocacy for its adoption spanning several decades [9]. The recent surge of decolonisation research calls special attention to the harm of colonialism to higher education. However, most research and movements hitherto focus primarily on humanities programmes, while the discussion within STEM remains limited, with even less within the biomedical sciences. Due to the global changes in the economic and political landscape, both patients and student cohorts in biomedical sciences encompass increasing cultural and ethnic diversity in the UK [10, 11]. To meet the diverse needs of both research and educational practice, the biomedical curricula need to develop and diversify to ensure that students acquire the knowledge, attitudes and skills necessary to become ethical and effective researchers. This is particularly important in the biomedical sciences, where research often translates into medical applications; consequently, scientific bias can lead to healthcare inequalities. For example, the vast majority of participants in genome-wide association studies (GWAS) are of European descent, leading to lower accuracy in clinical risk mapping for people of non-European descent [12, 13]. Therefore, it is imperative to raise awareness of the Eurocentric and colonial biases embedded in the discipline and increase exposure to a broad range of narratives and non-European paradigms.

Several reports of decolonising activities from science faculties have been published recently, including useful resources to support decolonial activities in chemistry [14-16]. There is also work being undertaken in many universities to decolonise the medical and dental curricula [17, 18]. But a clear framework for how to go about decolonising biomedical science programmes is currently lacking.

There has been extensive discourse about whether universities or curricula can truly be decolonised, particular from the position of a former coloniser, *metropolitan* institute. We acknowledge the problematic nature of decolonisation as a metaphor [19]; however, decolonising the curriculum has become the term of reference for a movement towards acknowledging colonial structures in our institutions, disciplines and pedagogies [1]. Use of the term *decolonising* implies a specific intent to explore and address the injustice and racism that are a legacy of colonialism; this is not adequately captured if the more comfortable, less divisive, label of *inclusive curriculum design* is applied. Much of what we are likely to achieve could more realistically be termed *diversifying* the curriculum, which we see as a step towards positive change with a decolonial mindset [20]. With this in mind, we aimed to evaluate our position on our decolonial journey within the biomedical sciences Schools at the University of Bristol. Our research questions were:

- What is our staff and students’ understanding of and attitudes towards decolonising and diversifying the curriculum?
- How inclusive is our current teaching practice?

Given the limited resources currently available in the literature, we sought to take an inductive approach by exploring the experience of students and staff members who are on the front lines of learning and teaching through the current curriculum. Thus, surveys and focus groups have been adopted as appropriate research methods. Using this mixed methods approach, we aimed to distil the outcomes into a framework describing characteristics of a decolonised curriculum. We propose that this framework can be used to develop a diverse and inclusive curriculum, placed within the appropriate historical context, and advocating for the equal and fair representation of all.

## Methods

### Survey data collection

Survey data were collected using Jisc Online Surveys. The survey was open from 23rd February to 23rd March 2022 and advertised to teaching staff and undergraduate students within the three biomedical sciences Schools within the University of Bristol, UK: Schools of Biochemistry, Cellular & Molecular Medicine (CMM) and Physiology, Pharmacology & Neuroscience (PPN). Survey questions are listed in the Supplementary Information.

Quantitative data were analysed using SPSS. Five-point scale questions were analysed non-parametrically using Mann-Whitney U tests for two-way comparisons and Kruksal-Wallis tests for more than two groups. Where there were significant differences (p<0.05) from Kruksal-Wallis tests, pairwise comparison was performed using Dunn post-hoc analysis with Bonferroni adjustment. Binary data were analysed using a Pearson Chi-square test, with post-hoc pairwise analysis using the Bonferroni adjustment for multiple comparisons.

### Focus groups

Four online focus groups were conducted: three with students across all year groups and one with staff members, all from biomedical sciences Schools. Participants were asked a series of six questions, all designed to avoid any perception of leading or threatening, while focusing on the participants’ understanding and experiences of potential colonialism in the biomedical sciences curricula. Detailed questions are shown as follows:

1. What do you understand by decolonising the curriculum? /What does a decolonised curriculum look like to you?
2. Do you think decolonising the curriculum is important? Why?
3. What do you understand by inclusive teaching practices?
4. What examples have you come across of inclusive teaching practices?
5. What examples have you come across of non-inclusive teaching practices?
6. Within your course, what could we improve in terms of inclusivity?

Automated transcripts from the online meeting platform were manually checked for accuracy. As the focus groups collected sensitive data from a small group of people, pseudonyms were used throughout to protect the confidentiality and privacy of the participants. We were cautious of power and voice issues intrinsic to (re)naming, especially its psychological and sociocultural implications to the participants and hence the research output [21]. Therefore, all the participants were invited to choose their own pseudonyms rather than being allocated one by the researchers.

The transcripts and responses to the open-ended questions from the surveys were analysed using an inductive approach with iterative strategies using NVivo 12 software (v. 12.7.0) and Microsoft Excel (v. 16.68) [22-24]. All qualitative data were first open-coded and then examined and axial-coded into categories that draw cogent connections between the open codes. The open and axial-coding processes were not linear but iterative, comprising several rounds of sorting and sifting so that the researchers could better understand the dimensions of the data and explore their relationship to it [25].

During theme development, ideas that appeared across the categories were identified as potential themes. Researchers then organised and assessed the prevalence and significance of the supporting evidence for that theme. All the final themes are well supported across both sources of data, i.e., focus groups and free-text responses from the surveys, and demonstrate systematic relations across the participants. Supplementary Table 1 displays the codebook containing categories, sub-themes, and themes of the 3Rs framework.

### Ethical review and open access

This study was reviewed and approved by the Faculties of Life Sciences and Science Research Ethics Committee at the University of Bristol in February 2022 (project number 9913). Participants completed an online consent form, agreeing to the open access use of their anonymised data, explaining how data would be stored, clarifying that their participation was voluntary and up to which point they could withdraw their data. Anonymised transcripts and survey responses are available via the Open Science Framework (https://osf.io/ps4ua/).

## Results & Discussion

### Staff are more familiar with decolonisation work than students

We surveyed teaching staff and undergraduate students on their understanding of and attitudes towards decolonising and diversifying the curriculum. Responses and analyses of the quantitative data are shown in Supplementary Tables 2-14. Staff (N=71) were significantly more familiar with decolonising than students (N=121): the median response from staff was ‘very aware’ whilst the median from students was ‘some awareness’; 20% of students had never heard of decolonising the curriculum (Figure 1A, Supp Table 2). This is consistent with a survey at Kingston University, where students’ understanding of decolonising was also quite low [16].

**Figure 1.**
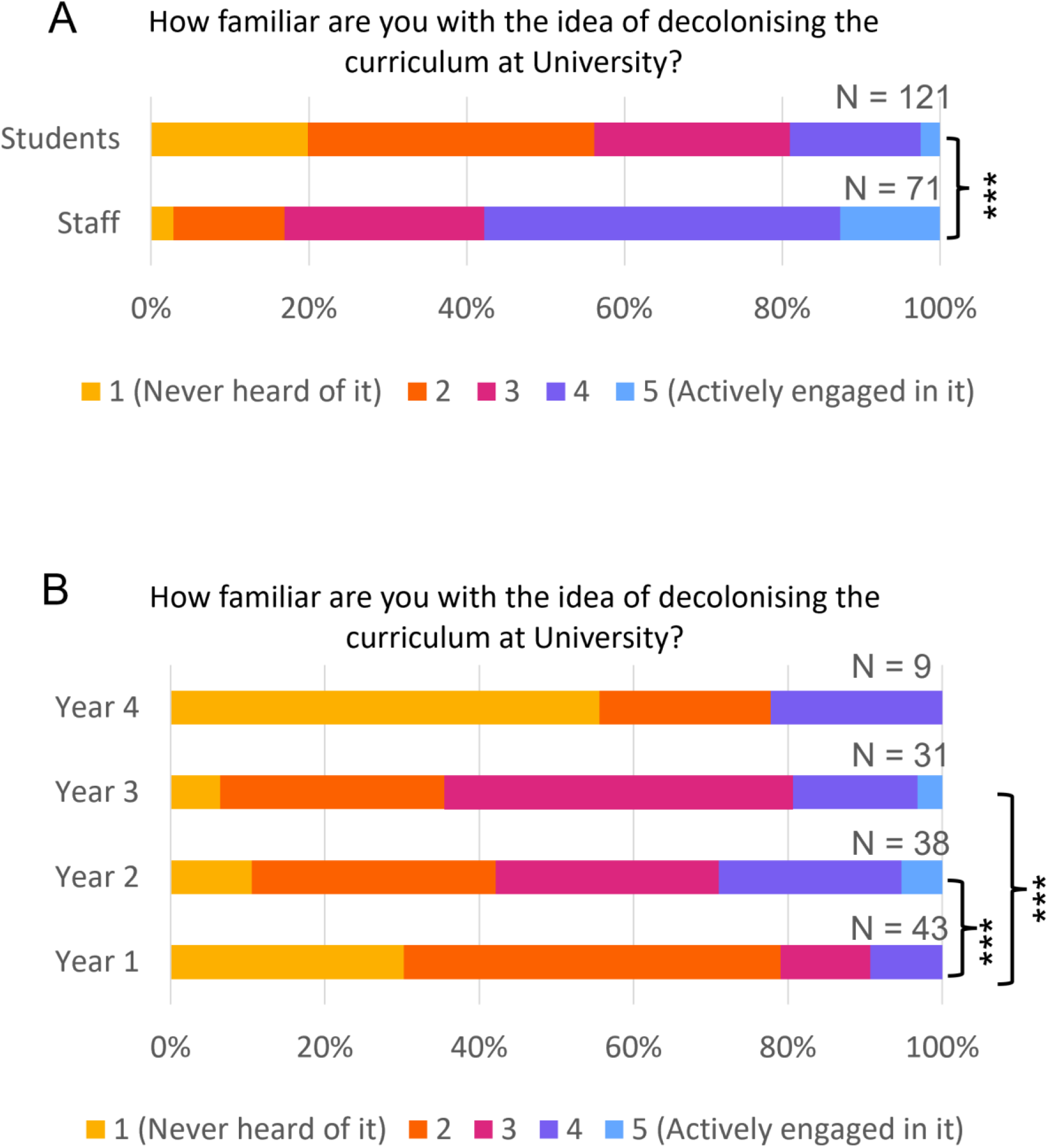
Familiarity with decolonising the curriculum. A) Comparison of responses from staff and students to the question ‘How familiar are you with the idea of decolonising the curriculum at University?’ 1 – never heard of it; 2 – some awareness; 3 – moderately aware; 4 – very aware; 5 – actively engaged in it. *** p<0.001 (Mann-Whitney U test). B) Comparison of responses from students in different year groups to the question ‘How familiar are you with the idea of decolonising the curriculum at University?’ *** p<0.001 (Kruskal-Wallis test, with Dunn-Bonferroni post-hoc pairwise comparisons).

On the student survey we collected personal information such as year of study, gender, ethnicity, sexual orientation, disability status and religion, enabling us to perform a demographic analysis (this was not possible on the staff survey due to the likelihood of deanonymising the data). There was a significant difference in familiarity with decolonising between different year groups (p<0.001), with Year 2 and 3 students more familiar than Year 1. This difference could reflect activities undertaken in the biomedical sciences Schools aimed at the Year 2 and Year 3 cohorts, such as: the addition of teaching and learning materials around decolonising the curriculum into a mandatory course for all Year 2 students, employment opportunities for students working on projects in this area, and information shared with students in newsletters. Surprisingly, this trend did not persist to Year 4 students, who appeared to be less familiar, although the number of students in this group is small and the difference here is not significant (Figure 1B; Supp Table 2). There were no significant differences in familiarity with decolonising activities between students of different gender, ethnicity, sexual orientation or religion. However, students who declared a disability were more familiar with decolonising the curriculum than those who did not (p=0.032; Supp Table 2).

### Importance of decolonising highlights gendered differences in students

Both staff and students thought it was important to decolonise the curriculum, with both groups giving a median response of 4 out of 5 (Figure 2A; Supp Table 3). Much of the recent drive to decolonise the curriculum has come from student campaigns [2, 26], so we were surprised that there was equal importance perceived by staff and students in our study. Female students thought it was more important (median 5 out of 5) than male students (median 4 out of 5; p<0.001; Figure 2B; Supp Table 3). Perhaps this reflects that women are more likely to identify as being marginalised in STEM subjects. However, there was no significant difference in perceived importance from students of different ethnicities, or between different sexual orientations, disability status or religion (Supp Table 3). Staff data were not analysed by demographic due to the risk of deanonymisation, so we cannot determine if these gender differences were also present in staff perceptions.

**Figure 2.**
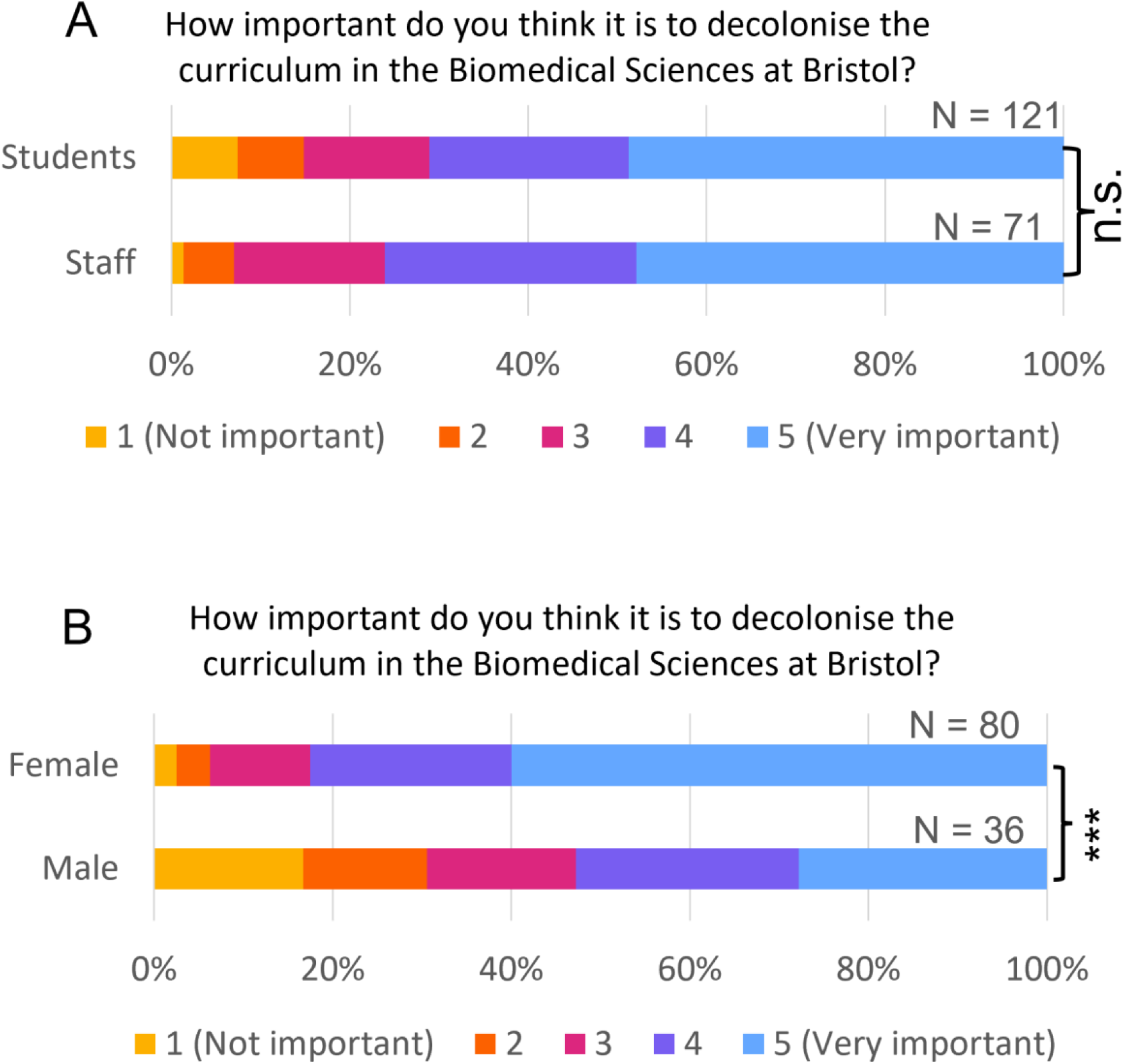
Importance of decolonising the curriculum. A) Comparison of responses from staff and students to the question to the question ‘How important do you think it is to decolonise the curriculum in the Biomedical Sciences at Bristol?’ 1 – not important; 5 – very important. B) Comparison of responses from female and male students to the question to the question ‘How important do you think it is to decolonise the curriculum in the Biomedical Sciences at Bristol?’ 1 – not important; 5 – very important. *** p<0.001 (Mann-Whitney U test).

### Students from minoritised ethnic groups feel less represented by the curriculum than white students

White students felt significantly more represented by the science and scientists they learnt about in the curriculum compared to Asian or black students, or those who gave their ethnicity as ‘not listed’ (Figure 3; Supp Table 8). The majority of the black students felt ‘not at all’ represented, whilst the median for Asian and mixed-race students was only 2 out of 5, compared to 4 out of 5 for white students. White students and staff are in the majority in our Schools, and the lack of a feeling of representation in the curriculum for non-white students likely reflects the Eurocentric nature of the field and the curriculum.

**Figure 3.**
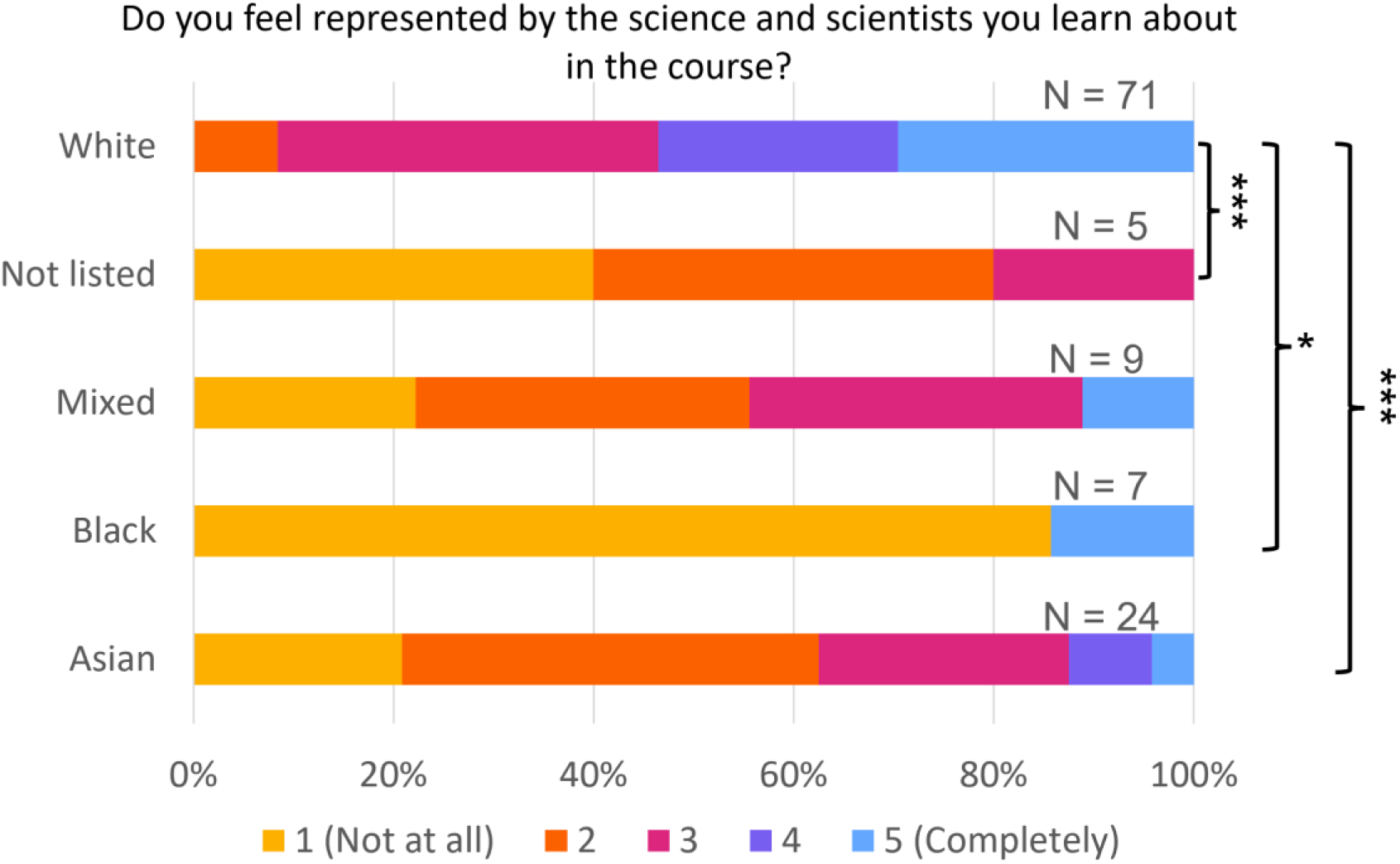
Representation in the curriculum. Comparison of responses from students of different ethnicity groups to the question ‘Do you feel represented by the science and scientists you learn about in the course?’ 1 – not at all; 5 – completely. * p<0.05; *** p<0.001 (Kruskal-Wallis test, with Dunn-Bonferroni post-hoc pairwise comparisons).

### Staff are more aware of decolonising activities in higher education than students

Digging deeper into the awareness of staff and students about decolonising activities, we asked whether participants were aware of activities currently taking place to decolonise the curriculum at different levels: UK universities in general; the University of Bristol (UoB); the Faculty of Life Science; their School or their specific modules (known as ‘units’ at UoB). Staff were more aware than students of decolonising activities, and this was significant at all levels within the University (Figure 4A; Supp Table 9). Staff were less aware of activities at UK universities in general compared to activities within the institution. This may reflect the active attempts of staff engaged in decolonising activities to share their work with colleagues at a local level. Students were more aware of activities at UoB than any other category, but less aware of decolonising work happening within their individual units (Figure 4A; Supp Table 10 and 11).

**Figure 4.**
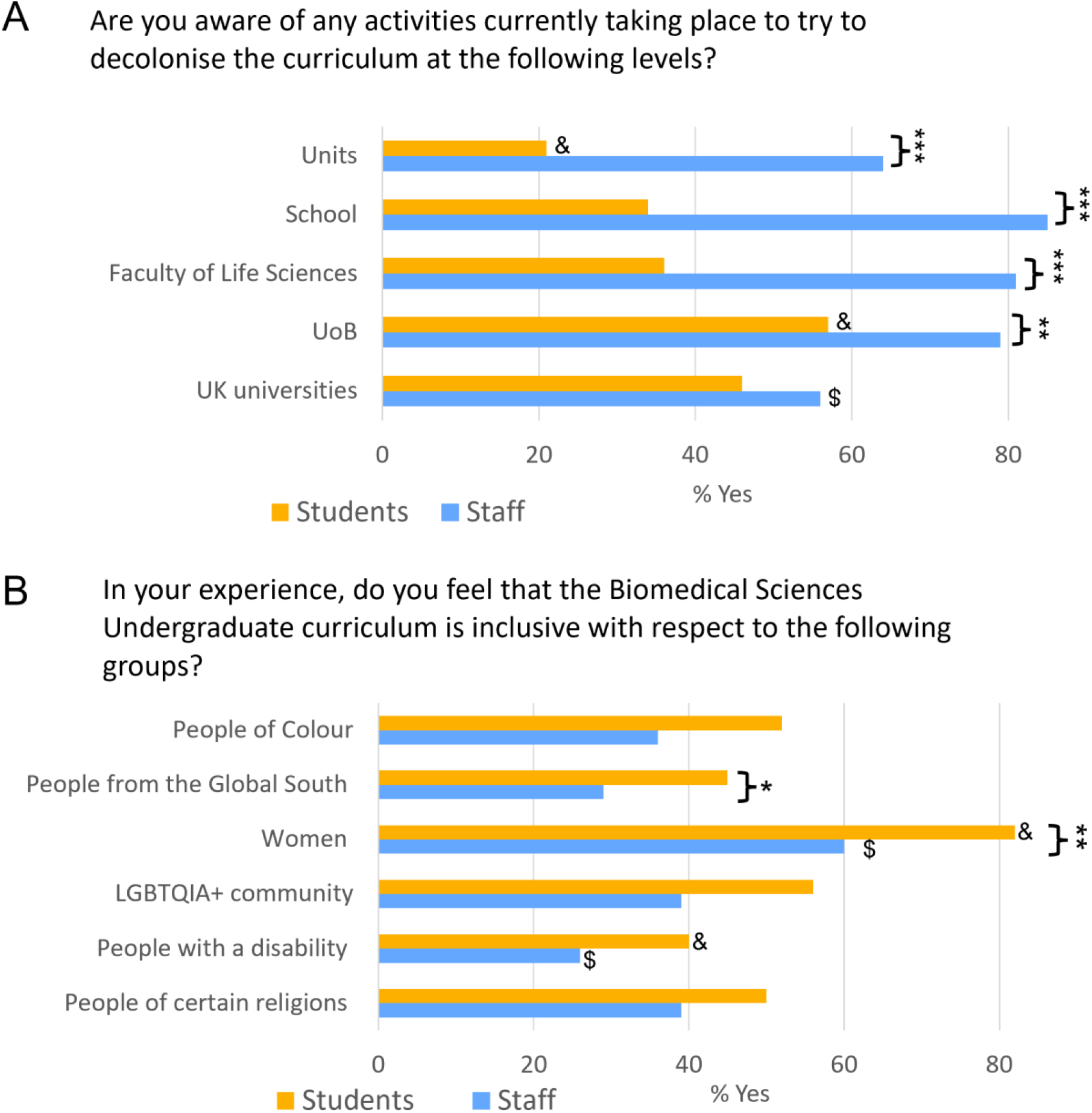
Awareness and Inclusivity. A) Responses to the question ‘Are you aware of any activities currently taking place to try to decolonise the curriculum at the following levels?’ shown as % of ‘Yes’ responses. B) Responses to the question ‘In your experience, do you feel that the Biomedical Sciences undergraduate curriculum is inclusive with respect to the following groups?’ shown as % of ‘Yes’ responses. * p<0.05, ** p<0.01, *** p<0.001 comparing staff and student responses (Pearson’s Chi-squared tests). $ p<0.05 comparing staff data across different categories; & p<0.05 comparing student data across different categories (Pearson’s Chi-squared tests and post-hoc test with Bonferroni correction).

### The curriculum is not viewed as highly inclusive, and staff feel it is less inclusive than students

Most participants thought the curriculum is not currently inclusive to marginalised groups. Compared to students, staff felt the curriculum was less inclusive, and this difference was significant for inclusion towards women and towards people from the Global South. Among both staff and students, more participants felt the curriculum was inclusive to women than to other marginalised groups, and fewest participants felt it was inclusive to people with a disability (Figure 4B; Supp Tables 12-14).

### Three themes for decolonising and diversifying the curriculum: rediscovery, representation and readiness

Thematic analysis of the focus group transcripts and free-text responses on the survey revealed three main themes:

- Rediscovery
- Representation
- Readiness

### Theme 1 - Rediscovery: alternative canons of knowledge

Epistemic injustice, i.e., the wrongs stemming from bias and inequity in knowledge generation, circulation and application [27, 28] emerged as a key topic in all focus groups. For instance, when discussing the motivations for decolonising the curriculum, one student participant commented:

> *I’ve realised studying science for, like most of my life, most of the scientists who [we] study about are white. So maybe it’s also crediting other people for their achievements because you usually don’t hear about them as much*.

Similar observations from staff members were found in the survey. The discussion here will first briefly synthesise and analyse the participants’ reflections on the Eurocentrism or “whiteness” of the curriculum, followed by an introduction to the initiatives that participants adopted aiming to redress epistemic injustice; this will lead to the closing discussion on the challenges that participants have confronted during the decolonisation process.

#### 1.1 Sources of information or Informants?

Epistemic injustice is a multi-faceted issue, which manifests in the dominant knowledge system and is strengthened by everyday knowledge practices. For example, as Emma ^1^ reflected:

> *We learn all about the experiments and data collected from just like British scientists. I’m sure there’s plenty of room for scientists from other places but we just don’t know about*.

This is in line with the observations from other participants about race and gender bias and hero-worshipping concerning scientific discovery. As Maria strongly argued:

> *I think we learn about the end point of how we got to the final conclusion, but so many people, maybe even 10, 20 even more contributed to that discovery being made that we’re not necessarily made aware of. So it’s not that one person came up with what the structure of DNA is actually, it was a lot of people and a lot of research, lots of small, tiny discoveries that led to the final big work being made*.

These observations mirror the testimonial injustice entrenched in academic research and curriculum, where knowledge held by people from marginalised groups, e.g., indigenous communities or women of colour, is systematically discredited and their status as knowers is prejudicially rejected [27]. As the participant’s reflection suggested, too often those belonging to marginalised groups are objectified as *sources of information* (much like arboreal growth rings, from which the inquirer can glean information) rather than *informants* (epistemic agents who convey information) [29]. While they both contribute to knowledge generation, the concept of knowledge is only tied to the latter.

Consequently, knowers from minoritised backgrounds face unreasonable obstacles when putting their knowledge in the public domain (see for example [30]). This inevitably leads to the underrepresentation of marginalised groups in the collective body of knowledge, which in turn may further impede their ability to understand or communicate the experiences effectively. At the aggregate level, any knowledge system that differs from the dominant one risks being relegated to an inferior position, despite being efficacious to the community it serves (see [31] for an example). Meanwhile, it unfairly grants advantages to those who are represented as having the power to structure collective social understandings. A consequence of such privileges is, as Bhambra *et al*. argue, that the current curriculum remains primarily “governed by the West for the West” [32].

#### 1.2 Retrieve Marginalised Knowledges

Epistemic injustice is widely acknowledged and underscored as a substantive issue by the participants. More importantly, they endeavoured to rectify it by taking the initiative themselves. For instance, many staff members reported that they have tried to diversify the reading list by including “proactive acknowledgement of contributions from scientists from minority groups”, “examples of literature from the Global South and a diversity of authors”, “[r]ecognition of the importance of indigenous cultures,” and “active presentation of alternate perspectives from other cultures/parts of the world; acknowledgement of diverse historical contribution to modern Life Sciences; acknowledgement of when narrow perspective has disadvantaged minorities in the past”.

Students also showed appreciation for the conscious shift towards addressing Eurocentrism in the curriculum at the teacher and institutional levels. As one student reflected on their lesson about the Tuskegee syphilis study [33]:

> *One of the lectures that stood out to me is that some of the experiments, like some of the ways that the scientists gained the data, was really really unethical (…) I think the inclusion of that with one of the lecturers was just so memorable (…) I didn’t, we just don’t really hear about it (…) I think it was really interesting (…) the idea that all these diseases one hundred years ago, they got to the stage magically, but there wasn’t actually a reasoning or, you know, how they got it, why they got it. And realistically, it probably was in underprivileged groups, people who didn’t have money (…) it’s quite interesting and maybe that should be included a little bit more in the lectures*.

Similarly, another student responded positively to the changes in the perceived learning environment, which they think “has been a real shift between last year and this year and the way that the curriculum has kind of approached it (…) they [staff] have made the effort to include more case studies and more diverse opinions and diverse people”. Interestingly, one participant noted assessment or coursework as an opportunity for students to question the implicit biases within the curriculum. As Ruby noted, “including these critical points in the essays”, namely the critical analysis of clinical and scientific issues related to ethnic minorities, “get you high marks (…) So I think you just gotta include a wide range of viewpoints”.

As shown above, staff participants took explicit efforts to include silenced or marginalised knowledge and researchers, attempting to address the power imbalances embedded in global knowledge production. Moreover, some participants have increasingly grown aware of the historical and anthropological underpinnings of the biomedical curriculum. It is this awareness that renders them more inclined to acknowledge, and where possible reverse, the policies and practices that uphold colonial legacy.

#### 1.3 Awareness, pedagogical practices & challenges

It is noteworthy that despite their commitment to epistemic decolonisation, many participants reported they faced challenges when translating its policy and guidelines into teaching and learning practices. As Cora, a staff member, noted:

> *[W]hen we give reading lists to students (…) particularly if it’s things that were experiments that were done many years ago, like in the fifties and sixties, they were often done by white male[s] (…) for some things we don’t have examples where we can broaden the older literature (…) it’s quite tricky*.

In addition to the limited access to the resources or networks of information about decolonisation, some staff also showed hesitance because of time or the additional workload, stating concerns such as “I’m not sure how I would fit it in to an already busy student timetable and with my heavy workload”. However, they appear willing to increase their efforts if provided with additional guidance and support.

Despite the challenges identified above appearing pervasively in the focus groups, these are rarely discussed in the existing literature. While much scholarly attention has focused on proposing guidance on what decolonisation efforts should be taken, little emphasis has been placed on the potential challenges or dilemmas met while implementing the efforts, and even less on the concrete challenges contextualised in the biomedical sciences. The process of decolonising or undoing colonialism in knowledge generation is a multifaceted and sophisticated one, inevitably involving uncomfortable truths. This is particularly relevant to attempts that aim to dismantle the unequal hierarchies which root much of the Western medical establishment [18]. The lack of discussion about these challenges risks reducing epistemic decolonisation to a simple task, while in reality, it requires substantive efforts. The findings also indicated that more resources and institutional support to academic staff can be pivotal to enabling the translation from awareness to pedagogical practices.

### Theme 2 - Representation: towards a more comprehensive understanding

The second theme focuses on the endeavours and remaining problems towards embedding diversity and inclusivity in the curriculum. As discussed in sub-theme 2.1, staff adopted effective practices to diversify the examples in the taught curriculum. Meanwhile, implicit bias still plagued the selection of topics, hampering an equity-oriented view of global biomedical research. Sub-theme 2.2 looks at race-ethnicity and gender representation in the faculty and its implication for diversifying the curriculum.

#### 2.1 Challenge the “norm”

During the focus groups, endeavours in diversifying the curriculum were reported from both staff and students’ experience in recent times. For example, as a student recalled:

> *I have an example, actually, I quite appreciated this. I was doing [deleted for confidentiality purposes] lecture and were learning about autoimmune diseases and one of them we learnt about was lupus. The lecture provided pictures of common symptoms. It was like a butterfly rash on the face. And she provided pictures of someone who had lighter skin, but also someone who had darker skin. You can see that there’s a clear difference in how the rash presents. And if you only saw the picture of the lighter skin and you saw that same rash on a dark-skinned person, you might not even know that they had lupus because it’s not as prominent. And that’s like another problem, I guess, the diseases or disorders on darker skin that they present a lot differently. But then a lot of studies are done on lighter skin, so they don’t know about what it looks like on darker skin, which can lead to, I guess, institutional racism because you would misdiagnose someone because you don’t know how it presents on their skin tone. So, I think that’s really important. I really appreciated that*.

A staff participant has also discussed how they tried to provide “[e]ven more diverse examples of factors influencing health problems around the globe”. According to them, “[biomedical studies] serve a wide population. The curriculum should reflect that by including different skin tones and genders instead of having sort of a norm”. Similarly, other considerations have also been in place questioning the idea of the “norm” patients, who are often portrayed as white heterosexual males.

In contrast to the increasingly diversified taught curriculum, the null curriculum, namely what is not taught, still conveys mixed messages. For example, many students mentioned “[a] lot of diseases can be more prevalent in other ethnic groups”, which however were never mentioned in the curriculum. As Gebrial pointed out: “Any curriculum must, by definition, exclude - the question is what is excluded and why” [18]. It is beyond the scope of this research to explore the full extent of the null curriculum in biomedical sciences and the reason for it. However, the information gleaned from the study sheds light on participants’ critical awareness of specific excluded topics and their demand for knowing about them, which warrants further interrogation of its implication.

#### 2.2 (Missing) Staff diversity matters

There has been an increasing call for staff diversity in UK higher education [34, 35]. Despite this, biomedical sciences, a discipline historically dominated by white European men, shows minimal progress in addressing race-ethnic and gender disparities during career progression [36, 37]. According to the student participants, the lack of diversity among faculty members could convey mixed messages, contradictory to the diversity tenets. For instance, as a student recalled:

> *I can say from my experience in biochemistry that I think I’ve only ever been taught by one lecturer, a person of colour in some of the 50 lecturers that have taught me in the four years that I’ve been at uni*.

In addition, students also discussed the impact of the lack of diverse staff members on their learning quality. As they observed:

> *I think might be thinking of more diverse hiring practices as well of the lecturers themselves because I think there is a very, very specific type of lecturer and of course, that is kind of a white man. And for one of my units, I’ve only been taught by white men. And it’s because I know obviously there are lots of women scientists coming into the field, so maybe when hiring* … *maybe be more aware of, you know, the students do want to be taught maybe even by more diverse people? And because I do think as well, it gives them a different approach to it, as women lecturers have maybe a different approach to the subject than my male lecturers* … *And I just think having a team of staff that do reflect a wider range of people is helpful, especially like the admin team as well. Sometimes you want to, you know, you want to approach someone you think you’re going to be comfortable with*.

As shown, students expect to have more diverse representation among faculty members. Numerous studies consistently report that a diverse faculty means not only diverse lived experience embedded in course content, but also scholarly interests, teaching methods, student support and unofficial norms, behaviours and values [38]. In other words, the more diverse the teaching staff, the greater the diversity of both the ‘formal’ curriculum (that which is actively taught) and the ‘hidden’ curriculum (the learning and values students are expected to pick up at university, without being explicitly taught [39]). The following reflection from a staff member can effectively demonstrate how a teacher’s lived experience of being a minority can drive them to better support their students who are in a similar position:

> *So I thought, you know, let’s change some of the names here to make it more inclusive* … *when I was a kid, they had this children’s television programme and at the end the lady who hosted it, she had this magic mirror that she used to look through it. It was just a circle in a hand, and she used to look through and say, who I can see? Oh, I can see Jenny and Peter and Sally! And then when you’re a kid, you’re like sitting around the TV thinking she’s just going to say my name. And I didn’t have a very typical Anglo-Saxon name* … *So [my name] never came out for that thing. When I was about three or four, that really stuck with me. That all, you know, you’d go to those shops where you’d buy something with the names, and my name was never there because it wasn’t a typical name. So, you know, I really had this strong urge that I had to change these names because it just really annoyed me throughout my whole childhood that I didn’t felt like my name was represented. And I can only imagine other people feel it even more strongly as I am [after all] white* … *So yeah, that’s one of the things I’ve been doing*.

In conclusion, embedding diversity and inclusivity in curricula is a complex and multi-faceted task, especially when null and hidden curricula are considered. As discussed, the participants recognised the need to encompass diverse attitudes and inclusive practices in the curriculum. This is supported by the quantitative data showing that students from minoritised ethnic groups felt less represented in the curriculum (Figure 3). However, the implicit biases of the discipline and individuals may impede progress. Raising awareness of these issues through department research could be an effective way to combat the biases. Furthermore, increased representation from minority groups among staff could be the key to building a more diverse culture. Though some progress has been made in recent years to increase ethnic diversity amongst staff at UK higher education institutions, minority ethnic staff continue to be underrepresented [40].

### Theme 3 - Readiness: students as agents of change

Students are pioneers in the higher education decolonisation movements. On March 9, 2015, Chumani Maxwele, a student from the University of Cape Town (UCT) in South Africa, threw containers of faeces against a statue of Cecil John Rhodes on the university’s campus [2]. This event catalysed the formation of #RhodesMustFall, a student-led movement that called into question the colonial legacy of Cecil Rhodes. This statue at UCT was removed from campus within a month. However, it also drove a proliferation of campaigns in South Africa - which later spread internationally - about decolonisation and structural change in universities. Examples include the #RhodesMustFall campaign at the University of Oxford and “Why is my curriculum white?” at University College London [26]. The demands of campaigns vary, but they were all initiated and organised by the students.

Student participants in this project also conveyed a strong voice, with most students surveyed being strongly invested in decolonising the curriculum (Figure 2A). They questioned the very nature of knowledge in the curriculum, as shown in previous sections, and demanded epistemic pluralism, striving to act as agents of change. As shown in the following quotes, they were aware of the complexities of real-world problems and believed only diverse theories, methods, and perspectives could equip them with the competence to overcome the challenges.

> *[F]or the longer term, considering that, well, maybe the majority of us going to be scientists, maybe communicate with the public and lots of different things. I think it’s really important just having a good awareness of how the content of our course actually does affect a lot of people. And it’s not like the whole public is just like one set of [homogenous] people*.
>
> *Well, I think it can be really important because sometimes smaller groups can be affected by the disease, and maybe there might not be as much finding about that. So it’s very good to raise awareness to the people who are going to be the scientists of the next generation and on where they could potentially want to focus research on areas that are maybe not as explored as others*.
>
> *Because I guess the knowledge that we gain from the university can be really impactful into the industries and jobs that we’re going to in the future*.
>
> *If science isn’t diverse, it’s not actually researching our society* … *I think we have to research what’s actually happening at the time*.

Some staff members also underscored the importance of including students’ voices in curriculum development and programme governance structures. “Student engagement/feedback in [curriculum] design” was listed as a key action for decolonising the curriculum in the survey results. This echoes wider recommendations on the vital importance of co-production, where students work in partnership with staff to develop strategies to address ethnicity degree awarding gaps at UK higher education institutions [40]. Additionally, one staff participant suggested that special consideration ought to be given to students from underrepresented groups. Previous studies surveying students’ learning experiences in UK universities report that students of colour may be alienated as their histories and ancestral narrative are often omitted from mainstream discourse [26, 41]. To confront the Eurocentrism in the curriculum and institutional racism, it is essential to listen to the voices of minority groups and empower them to challenge their preconceptions and act as agents of transformation.

## Conclusions

This paper details the findings of a project aiming to understand staff and student attitudes towards decolonising and diversifying the biomedical sciences curricula at the University of Bristol. Quantitative data collected from a relatively large sample of staff and students were combined with the qualitative data from focus groups and free text survey questions. This allowed us to glean a more in-depth understanding of the mechanism underlying the survey results [42]. Further, the mixed method design enabled us to triangulate the findings derived from various data sources, increasing the validity of their conclusion. It also added breadth and scope to the project by shedding light on different facets of staff and students’ understanding of and attitudes towards decolonising and diversifying the curriculum.

Quantitative data revealed that students and staff are equally invested in decolonising and diversifying the curriculum, but students are less aware of the current activities taking place to this end (Figures 1 & 2). Among students, women felt this was more important than men, whilst students from minority ethnic groups felt less represented within the curriculum than white students, highlighting different perspectives of minoritised groups (Figure 3). Thematic analysis of the qualitative data using an inductive approach discovered three main themes, namely, rediscovery, representation and readiness.

The first theme, centred around epistemic injustice, concerns the power imbalance in the generation, circulation and application of ‘legitimate’ knowledge. It reveals the participants’ endeavours to rediscover the knowledge and scientists discredited by epistemic colonialism. Furthermore, it offers insights into the challenges the participants confronted while redressing this epistemic injustice.

The second theme emphasises embedding diversity and inclusivity in the ‘formal’ (explicit), ‘hidden’ (implicit) and ‘null’ (excluded) curriculum. The staff participants adopted effective strategies to challenge the traditional idea of the ‘norm patient’ and to address the diverse educational needs. Going forward, increased staff diversity could be the key to developing a culturally-responsive and equity-oriented biomedical science education.

The final theme reflects on the role of students’ voices in ongoing decolonisation initiatives. Across decolonisation movements globally, students have consistently pioneered, rendering the initiatives a ground-up feature. In this project, many student participants also expressed their enthusiasm for an equity and inclusivity-oriented curriculum. As the staff suggested, future decolonisation efforts will require empowering all students to challenge the status quo and act as agents of change.

We propose that these three themes can form a framework for a decolonised curriculum that could be useful for decolonial activities in the biomedical sciences and beyond. This is not only important for social justice and reparations for past colonial wrongs, but also imperative for ameliorating healthcare inequalities [13]. A recent largescale survey of students across the UK highlighted that an inclusive curriculum with diverse perspectives was considered important for academic rigour and a rounded view of the discipline. A perceived lack of inclusive content in the curriculum was associated with a poor sense of ‘belonging’ at university [43]. Bringing broader perspectives into the curriculum may therefore increase the rigour and relevance of the discipline and help students develop a better sense of belonging to the academic community.

Above all, this research reveals numerous opportunities for rectifying the colonial legacy embedded in the curriculum and most participants’ desire to do so. Furthermore, it attempted to shed light on the nuances in each individual’s implementation of decolonisation endeavours, especially the potential struggles and coping strategies. We hope this project has offered a safe space for staff and student participants to voice their concerns about the presence of colonialism in the institution and curriculum, and anticipate that the project’s findings will lead to further decolonisation dialogue and initiatives within UK higher education and beyond.

## Acknowledgements

We are very grateful for useful discussions with Dr Lara Lalemi, Prof Robin Shields, Prof Alvin Birdi, Prof Foluke Abedisi, Prof Leon Tikly, Dr Dave Lawson and Dr Celine Petitjean. We thank all the staff and students who responded to our survey and took part in focus groups. We are grateful for funding and support from the British Society of Immunology, Bristol Institute for Learning and Teaching (BILT), and the Faculty of Life Sciences Education Innovation Fund at the University of Bristol.

## Author contributions

TL planned and carried out focus groups, analysed the qualitative data and helped write the manuscript. LM performed statistical analysis of the quantitative data, under the supervision of AR. AR, CM, BB and AD applied for the funding, designed the survey instrument and wrote the manuscript. ZB and AM helped distribute the survey and helped write the manuscript.

## Footnotes

1 Pseudonyms are used for participants; see Methods for details.

## Notes

https://osf.io/ps4ua/

### Competing Interest Statement

The authors have declared no competing interest.

