## Supplementary Information for "A framework for decolonising and diversifying biomedical sciences curricula: rediscovery, representation and readiness"

**Supplementary Table 1.** Codebook with categories, sub-themes, and themes of the 3Rs framework. The left column comprises the categories that emerged from the raw data through open-coding and axial-coding; these categories are then clustered and refined to form the sub-themes in the middle column and, eventually, the themes in the right column. For Theme 3 Readiness, categories directly correspond to the themes in the right column.

| Categories | Sub-themes | Themes |
|---|---|---|
| Hero-worshipping: asymmetrical visibility in scientific discovery | *1.1 Sources of information or Informants?* | **Theme 1 Rediscovery**: alternative canons of knowledge |
| Credibility deficit: asymmetrical recognition in scientific discovery between Western researchers and the local community | | |
| Lack of acknowledgement of certain groups systematically excluded from knowledge production | | |
| Intend to enrich reading lists by including scientists from minority backgrounds | *1.2 Retrieve Marginalised Knowledges* | |
| Attempts to integrate historical unethical experiments as critical case studies in the curriculum | | |
| Promoting critical analysis of different ways of knowing and knowledge in essay-based assignments | | |
| Limited access to resources or networks of information on decolonisation efforts | *1.3 Awareness, Pedagogical Practices & Challenges* | |
| Time constraints of classes and competing priorities | | |
| Workload requirements exceeding individuals' capacity | | |
| Complexity in transitioning from awareness to pedagogical practices | | |
| Lack of training in addressing historical injustice | | |
| Incorporating compelling visual evidence, such as showcasing diseases on different skin tones | *2.1 Challenge the "Norm"* | **Theme 2 Representation**: towards a more comprehensive understanding |
| Promoting global awareness through case studies | | |
| Concerns about implicit bias in the null curriculum | | |
| Delayed diversification of staff: Concerns about the current predominance of white males among staff members and the expectation for broader representation | *2.2 (Missing) Staff Diversity Matters* | |
| Role modelling and inspiration: Lack of role modelling inspiration for students from underrepresented groups due to a lack of staff diversity | | |
| Curriculum content: Disproportionate staff representation leads to a Eurocentric scholarly focus in the curriculum content | | |
| Leadership diversity: The impact of lacking committee representation on decision-making and equitable outcomes | | |
| The complexities of real-world problems students face now and in the workplace. | | **Theme 3 Readiness:** students as agents of change |
| Enhancing the epistemic agency of the 'next generation' of scientists requires diverse theories, methods, and perspectives in the curriculum they receive. | | |
| Decolonised and diversified curriculum for fostering ethical and relevant scientific inquiry. | | |

**Supplementary Table 2.** Responses to the question 'How familiar are you with the idea of decolonising the curriculum at university?' 1 – never heard of it; 2 – some awareness; 3 – moderately aware; 4 – very aware; 5 – actively engaged in it. P values are from Mann-Whitney U tests for two-way comparisons and Kruskal-Wallis tests when there are more than two groups.

| **How familiar are you with the idea of decolonising the curriculum at university?** | | | | |
|---|---|---|---|---|
| **Demographic** | **Group** | **Median** | **N** | **P** |
| Staff vs Students | Students | 2 | 121 | <0.001 |
| | Staff | 4 | 71 | |
| Ethnicity (students) | Asian | 2 | 24 | 0.712 |
| | Black | 2 | 7 | |
| | Mixed | 2 | 9 | |
| | Not listed | 2 | 5 | |
| | White | 2 | 71 | |
| Gender (students) | Female | 2 | 80 | 0.943 |
| | Male | 2 | 36 | |
| Sexuality (students) | LGBTQIA+ | 2 | 45 | 0.197 |
| | Heterosexual | 2 | 64 | |
| Disability (students) | Disability | 3 | 19 | 0.032 |
| | No disability | 2 | 96 | |
| Religion (students) | Religious | 2 | 37 | 0.645 |
| | Not religious | 2 | 80 | |
| Year of study (students) | 1 | 2 | 43 | <0.001 |
| | 2 | 3 | 38 | |
| | 3 | 3 | 31 | |
| | 4 | 1 | 9 | |

**Supplementary Table 3.** Responses to the question 'How important do you think it is to decolonise the curriculum in the Biomedical Sciences at Bristol?' 1 – not important; 5 – very important. P values are from Mann-Whitney U tests for two-way comparisons and Kruskal-Wallis tests when there are more than two groups.

| How important do you think it is to decolonise the curriculum in the Biomedical Sciences at Bristol? | | | | |
|---|---|---|---|---|
| **Demographic** | **Group** | **Median** | **N** | **P** |
| Staff vs Students | Students | 4 | 121 | 0.640 |
| | Staff | 4 | 71 | |
| Ethnicity (students) | Asian | 4 | 24 | 0.68 |
| | Black | 5 | 7 | |
| | Mixed | 4 | 9 | |
| | Not listed | 5 | 5 | |
| | White | 5 | 71 | |
| Gender (students) | Female | 5 | 80 | <0.001 |
| | Male | 4 | 36 | |
| Sexuality (students) | LGBTQIA+ | 4 | 45 | 0.701 |
| | Heterosexual | 5 | 64 | |
| Disability (students) | Disability | 4 | 19 | 0.817 |
| | No disability | 5 | 96 | |
| Religion (students) | Religious | 4 | 37 | 0.268 |
| | Not religious | 5 | 80 | |
| Year of study (students) | 1 | 5 | 43 | 0.751 |
| | 2 | 4.5 | 38 | |
| | 3 | 4 | 31 | |
| | 4 | 4 | 9 | |

**Supplementary Table 4.** Responses to the question 'To what extent does your curriculum reflect the contribution of diverse cultures to the development of Biomedical Sciences?' 1 – not at all; 5 – completely. P values are from Mann-Whitney U tests for two-way comparisons and Kruskal-Wallis tests when there are more than two groups.

| To what extent does your curriculum reflect the contribution of diverse cultures to the development of Biomedical Sciences? | | | | |
|---|---|---|---|---|
| **Demographic** | **Group** | **Median** | **N** | **P** |
| Staff vs Students | Students | 3 | 121 | 0.679 |
| | Staff | 2 | 68 | |
| Ethnicity (students) | Asian | 2.5 | 24 | 0.137 |
| | Black | 1 | 7 | |
| | Mixed | 2 | 9 | |
| | Not listed | 3 | 5 | |
| | White | 3 | 71 | |
| Gender (students) | Female | 3 | 80 | 0.987 |
| | Male | 3 | 36 | |
| Sexuality (students) | LGBTQIA+ | 2 | 45 | 0.836 |
| | Heterosexual | 3 | 64 | |
| Disability (students) | Disability | 2 | 19 | 0.358 |
| | No disability | 3 | 96 | |
| Religion (students) | Religious | 3 | 37 | 0.156 |
| | Not religious | 2 | 80 | |
| Year of study (students) | 1 | 2 | 43 | 0.476 |
| | 2 | 3 | 38 | |
| | 3 | 3 | 31 | |
| | 4 | 3 | 9 | |

**Supplementary Table 5.** Responses to the question 'To what extent does your curriculum address the needs of diverse populations across the world?' 1 – not at all; 5 – completely. P values are from Mann-Whitney U tests for two-way comparisons and Kruskal-Wallis tests when there are more than two groups.

| To what extent does your curriculum address the needs of diverse populations across the world? | | | | |
|---|---|---|---|---|
| **Demographic** | **Group** | **Median** | **N** | **P** |
| Staff vs Students | Students | 3 | 121 | 0.994 |
| | Staff | 3 | 68 | |
| Ethnicity (students) | Asian | 3 | 24 | 0.672 |
| | Black | 2 | 7 | |
| | Mixed | 2 | 9 | |
| | Not listed | 3 | 5 | |
| | White | 3 | 71 | |
| Gender (students) | Female | 3 | 80 | 0.064 |
| | Male | 2 | 36 | |
| Sexuality (students) | LGBTQIA+ | 2 | 45 | 0.075 |
| | Heterosexual | 3 | 64 | |
| Disability (students) | Disability | 3 | 19 | 0.795 |
| | No disability | 3 | 96 | |
| Religion (students) | Religious | 3 | 37 | 0.358 |
| | Not religious | 3 | 80 | |
| Year of study (students) | 1 | 3 | 43 | 0.274 |
| | 2 | 3 | 38 | |
| | 3 | 2 | 31 | |
| | 4 | 3 | 9 | |

**Supplementary Table 6.** Responses to the question 'To what extent does your curriculum celebrate the contributions of people of colour and other minoritised groups to the development of biomedical sciences?' 1 – not at all; 5 – completely. P values are from Mann-Whitney U tests for two-way comparisons and Kruksal-Wallis tests when there are more than two groups.

| **To what extent does your curriculum celebrate the contributions of people of colour and other minoritised groups to the development of biomedical sciences?** | | | | |
|---|---|---|---|---|
| **Demographic** | **Group** | **Median** | **N** | **P** |
| Staff vs Students | Students | 2 | 121 | 0.580 |
| | Staff | 2 | 68 | |
| Ethnicity (students) | Asian | 2 | 24 | 0.844 |
| | Black | 1 | 7 | |
| | Mixed | 2 | 9 | |
| | Not listed | 2 | 5 | |
| | White | 2 | 71 | |
| Gender (students) | Female | 2 | 80 | 0.884 |
| | Male | 2 | 36 | |
| Sexuality (students) | LGBTQIA+ | 2 | 45 | 0.627 |
| | Heterosexual | 2 | 64 | |
| Disability (students) | Disability | 2 | 19 | 0.281 |
| | No disability | 2 | 96 | |
| Religion (students) | Religious | 2 | 37 | 0.280 |
| | Not religious | 2 | 80 | |
| Year of study (students) | 1 | 2 | 43 | 0.704 |
| | 2 | 2 | 38 | |
| | 3 | 2 | 31 | |
| | 4 | 2 | 9 | |

**Supplementary Table 7.** Responses to the question 'To what extent does your curriculum draw on examples relating to other cultures and people of colour to illustrate ideas/concepts?' 1 – not at all; 5 – completely. P values are from Mann-Whitney U tests for two-way comparisons and Kruskal-Wallis tests when there are more than two groups.

| **To what extent does your curriculum draw on examples relating to other cultures and people of colour to illustrate ideas/concepts?** | | | | |
|---|---|---|---|---|
| **Demographic** | **Group** | **Median** | **N** | **P** |
| Staff vs Students | Students | 2 | 121 | 0.077 |
| | Staff | 2 | 66 | |
| Ethnicity (students) | Asian | 2 | 24 | 0.099 |
| | Black | 1 | 7 | |
| | Mixed | 2 | 9 | |
| | Not listed | 2 | 5 | |
| | White | 2 | 71 | |
| Gender (students) | Female | 2 | 80 | 0.740 |
| | Male | 2 | 36 | |
| Sexuality (students) | LGBTQIA+ | 2 | 45 | 0.903 |
| | Heterosexual | 2 | 64 | |
| Disability (students) | Disability | 2 | 19 | 0.913 |
| | No disability | 2 | 96 | |
| Religion (students) | Religious | 2 | 37 | 0.382 |
| | Not religious | 2 | 80 | |
| Year of study (students) | 1 | 2 | 43 | 0.988 |
| | 2 | 2 | 38 | |
| | 3 | 2 | 31 | |
| | 4 | 2 | 9 | |

**Supplementary Table 8.** Student responses to the question 'Do you feel represented by the science and scientists you learn about in the course?' 1 – not at all; 5 – completely. P values are from Mann-Whitney U tests for two-way comparisons and Kruskal-Wallis tests when there are more than two groups.

| **Do you feel represented by the science and scientists you learn about in the course?** | | | | |
|---|---|---|---|---|
| **Demographic** | **Group** | **Median** | **N** | **P** |
| All Students | Students | 3 | 121 | - |
| Ethnicity (students) | Asian | 2 | 24 | <0.001 |
| | Black | 1 | 7 | |
| | Mixed | 2 | 9 | |
| | Not listed | 2 | 5 | |
| | White | 4 | 71 | |
| Gender (students) | Female | 3 | 80 | 0.156 |
| | Male | 3 | 36 | |
| Sexuality (students) | LGBTQIA+ | 3 | 45 | 0.877 |
| | Heterosexual | 3 | 64 | |
| Disability (students) | Disability | 3 | 19 | 0.737 |
| | No disability | 3 | 96 | |
| Religion (students) | Religious | 2 | 37 | 0.002 |
| | Not religious | 3 | 80 | |
| Year of study (students) | 1 | 3 | 43 | 0.117 |
| | 2 | 3 | 38 | |
| | 3 | 3 | 31 | |
| | 4 | 4 | 9 | |

**Supplementary Table 9.** Responses to the question ‘Are you aware of any activities currently taking place to try to decolonise the curriculum at [different levels]?’ Comparison of student and staff responses with Pearson’s chi square analysis.

| **Are you aware of any activities currently taking place to try to decolonise the curriculum at UK Universities?** | | | | |
|---|---|---|---|---|
| | **No** | **Yes** | **Total** | **Pearson’s chi square** |
| Staff | 28 | 36 | 64 | 0.216 |
| Students | 63 | 54 | 117 | |
| Total | 91 | 90 | 181 | |
| **Are you aware of any activities currently taking place to try to decolonise the curriculum at the University of Bristol?** | | | | |
| | **No** | **Yes** | **Total** | **Pearson’s chi square** |
| Staff | 14 | 54 | 68 | 0.002 |
| Students | 52 | 68 | 120 | |
| Total | 66 | 122 | 188 | |
| **Are you aware of any activities currently taking place to try to decolonise the curriculum in the Faculty of Life Sciences?** | | | | |
| | **No** | **Yes** | **Total** | **Pearson’s chi square** |
| Staff | 12 | 54 | 66 | <0.001 |
| Students | 75 | 42 | 117 | |
| Total | 87 | 96 | 183 | |
| **Are you aware of any activities currently taking place to try to decolonise the curriculum in your School?** | | | | |
| | **No** | **Yes** | **Total** | **Pearson’s chi square** |
| Staff | 11 | 60 | 71 | <0.001 |
| Students | 77 | 39 | 116 | |
| Total | 88 | 99 | 187 | |
| **Are you aware of any activities currently taking place to try to decolonise the curriculum in your units?** | | | | |
| | **No** | **Yes** | **Total** | **Pearson’s chi square** |
| Staff | 23 | 41 | 64 | <0.001 |
| Students | 90 | 24 | 114 | |
| Total | 113 | 65 | 178 | |

**Supplementary Table 10.** Student responses to the question 'Are you aware of any activities currently taking place to try to decolonise the curriculum at [different levels]?' Comparison of students' responses to different categories with Pearson's chi square analysis and post-hoc tests. Bonferroni adjusted alpha value = 0.005 for 10 comparisons.

| **Are you aware of any activities currently taking place to try to decolonise the curriculum at…?** | | | | | |
|---|---|---|---|---|---|
| | No | Yes | Total | P value | Post-hoc P value |
| **UK universities** | 63 | 54 | 117 | <0.001 | 0.0719 |
| **UoB** | 52 | 68 | 120 | | <0.001 |
| **Faculty of Life Sciences** | 75 | 42 | 117 | | 0.484 |
| **School** | 77 | 39 | 116 | | 0.194 |
| **Units** | 90 | 24 | 114 | | <0.001 |

**Supplementary Table 11.** Staff responses to the question 'Are you aware of any activities currently taking place to try to decolonise the curriculum at [different levels]?' Comparison of staff responses to different categories with Pearson's chi square analysis and post-hoc tests. Bonferroni adjusted alpha value = 0.005 for 10 comparisons.

| **Are you aware of any activities currently taking place to try to decolonise the curriculum at…?** | | | | | |
|---|---|---|---|---|---|
| | No | Yes | Total | P value | Post-hoc P value |
| **UK universities** | 28 | 36 | 64 | <0.001 | <0.001 |
| **UoB** | 14 | 54 | 68 | | 0.230 |
| **Faculty of Life Sciences** | 11 | 60 | 71 | | 0.0164 |
| **School** | 12 | 54 | 66 | | 0.0891 |
| **Units** | 23 | 41 | 64 | | 0.0574 |

**Supplementary Table 12**. Responses to the question 'In your experience, do you feel that the Biomedical Sciences undergraduate curriculum is inclusive with respect to the following groups?' Comparison of student and staff responses with Pearson's chi square analysis.

| **In your experience, do you feel that the Biomedical Sciences undergraduate curriculum is inclusive with respect to the following groups?** | | | | |
|---|---|---|---|---|
| | | **People of colour** | | |
| | **No** | **Yes** | **Total** | **Pearson's chi square** |
| Staff | 39 | 22 | 61 | 0.057 |
| Students | 56 | 60 | 116 | |
| Total | 95 | 82 | 177 | |
| | | **People from the Global South** | | |
| | **No** | **Yes** | **Total** | **Pearson's chi square** |
| Staff | 44 | 18 | 62 | 0.038 |
| Students | 63 | 52 | 115 | |
| Total | 107 | 70 | 177 | |
| | | **Women** | | |
| | **No** | **Yes** | **Total** | **Pearson's chi square** |
| Staff | 24 | 36 | 60 | 0.003 |
| Students | 21 | 94 | 115 | |
| Total | 45 | 130 | 175 | |
| | | **LGBTQIA+ community** | | |
| | **No** | **Yes** | **Total** | **Pearson's chi square** |
| Staff | 37 | 24 | 61 | 0.057 |
| Students | 51 | 64 | 115 | |
| Total | 88 | 88 | 176 | |
| | | **People with a disability** | | |
| | **No** | **Yes** | **Total** | **Pearson's chi square** |
| Staff | 45 | 16 | 61 | 0.096 |
| Students | 68 | 45 | 113 | |
| Total | 113 | 61 | 174 | |
| | | **People of certain religions** | | |
| | **No** | **Yes** | **Total** | **Pearson's chi square** |
| Staff | 36 | 23 | 59 | 0.197 |
| Students | 55 | 55 | 110 | |
| Total | 91 | 78 | 169 | |

**Supplementary Table 13.** Student responses to the question 'In your experience, do you feel that the Biomedical Sciences undergraduate curriculum is inclusive with respect to the following groups?' Comparison of students' responses to different categories with Pearson's chi square analysis and post-hoc tests. Bonferroni adjusted alpha value = 0.0042 for 12 comparisons.

| | No | Yes | Total | P value | Post-hoc P value |
|---|---|---|---|---|---|
| **People with a disability** | 68 | 45 | 113 | <0.001 | <0.001 |
| **People from the Global South** | 63 | 52 | 115 | | 0.0357 |
| **LGBTQIA+ community** | 51 | 64 | 115 | | 0.689 |
| **People of colour** | 56 | 60 | 116 | | 0.549 |
| **People with certain religious beliefs** | 55 | 55 | 110 | | 0.368 |
| **Women** | 21 | 94 | 115 | | <0.001 |

**Supplementary Table 14.** Staff responses to the question 'In your experience, do you feel that the Biomedical Sciences undergraduate curriculum is inclusive with respect to the following groups?' Comparison of staff responses to different categories with Pearson's chi square analysis and post-hoc tests. Bonferroni adjusted alpha value = 0.0042 for 12 comparisons.

| | No | Yes | Total | P value | Post-hoc P value |
|---|---|---|---|---|---|
| **People with a disability** | 45 | 45 | 113 | 0.003 | <0.001 |
| **People from the Global South** | 44 | 52 | 115 | | 0.0357 |
| **LGBTQIA+ community** | 37 | 64 | 115 | | 0.689 |
| **People of colour** | 39 | 60 | 116 | | 0.549 |
| **People with certain religious beliefs** | 36 | 55 | 110 | | 0.368 |
| **Women** | 24 | 36 | 115 | | <0.001 |

### Staff survey questions

*Part 1: demographic information*

Q1. Which is your home School?
Biochemistry
CMM
PPN

Q2. What is your academic career pathway[1]?
Pathway 1
Pathway 2
Pathway 3
Prefer not to say

Q3. Do you consider yourself to be a member of a minoritised group?
Yes
No
Prefer not to say

Q3a. If yes, please tell us which minoritised group or groups you belong to, if you feel comfortable doing so.

*Part 2: decolonising the curriculum*

Q4. How familiar are you with the idea of decolonising the curriculum at university?
Never heard of it
Some awareness
Moderately aware
Very aware
Actively engaged in it

*Decolonisation is broadly about confronting how colonialism, Eurocentrism, and racism have shaped our world. Decolonisation of the curriculum recognises these issues in the science we teach and aims to include a range of perspectives of people who have been overlooked, so as to present a truer picture of our discipline so that it can be more responsive to the needs of all.*

Q5. Are you aware of any activities currently taking place to try to decolonise the curriculum at the following levels?

| | Yes | No |
|---|---|---|
| UK universities | | |
| University of Bristol | | |
| Faculty of Life Sciences | | |
| My School | | |
| My units | | |

Q6. How important do you think it is to decolonise the curriculum in the Biomedical Sciences at the University of Bristol?
Not important 1 2 3 4 5 very important

Q7. In your experience, to what extent does the curriculum within the Biomedical Schools (Biochemistry, CMM, or PPN) reflect the contribution of diverse cultures to the development of biomedical sciences?
Not at all 1 2 3 4 5 Completely

Q8. In your experience, to what extent does the curriculum within the Biomedical Schools address the needs of diverse populations around the world?
Not at all 1 2 3 4 5 Completely

---

[1] There are three academic career pathways at the University of Bristol: Pathway 1 (research and teaching focused); Pathway 2 (research focused) and Pathway 3 (teaching and scholarship focused)

Q9. In your experience, to what extent does the curriculum within the Biomedical Schools celebrate the contributions of people of colour and other minoritised groups to the development of biomedical sciences?
Not at all 1 2 3 4 5 Completely

Q10. In your experience, to what extent does the curriculum within the Biomedical Schools draw on examples relating to other cultures and people of colour to illustrate ideas/ concepts?
Not at all 1 2 3 4 5 Completely

Q11. What three things would you expect to see in a decolonised curriculum? [Max 280 characters]

*Part 3 – Equality, Diversity and Inclusion in the Biomedical Sciences curriculum*
Q12 Do you feel all students are represented by the science and scientists that students learn about in the course?
Not at all 1 2 3 4 5 Completely

Q13. In your experience, do you feel that the Biomedical Sciences curriculum (School of Biochemistry, CMM or PPN) is inclusive with respect to the following groups?

| | Yes | No |
|---|---|---|
| People of colour | | |
| People from the Global South | | |
| Women | | |
| LGBTQIA community | | |
| People with a disability | | |
| People of certain religions | | |
| Not listed (please describe) _______________ | | |

Q14. Do you feel comfortable discussing issues of inclusivity, bias or inequality with students in the courses on which you teach?
- Very comfortable
- Somewhat comfortable
- Neither comfortable nor uncomfortable
- Somewhat uncomfortable
- Very uncomfortable
- Not aware of any

Q15. If you have experienced any non-inclusive practice (such as bias or inequality) at the university, at which level of the organisation was it? Select all that apply:
- Not experienced any non-inclusive practice
- Taught units
- Wider curriculum
- Personal tutoring
- University infrastructure or senior management
- Other_____________

Q16. If you are made aware of any non-inclusive practice (such as bias or inequality) in the courses in your school, would you feel comfortable raising it?
- Very comfortable
- Somewhat comfortable
- Neither comfortable nor uncomfortable
- Somewhat uncomfortable
- Very uncomfortable

Q17. To whom would you feel it is best to raise these issues if you had them? *Select all answers which apply.*

- Individual lecturers
- Unit Director
- School Education Director
- Faculty Education Director
- Head of School
- School Equality, Diversity and Inclusion Committee
- Other_____________

*Part 4 - Final comments*
Q18. Do you have any other comments relating to decolonising and diversifying the curriculum?

Q19. Would you be interested in working with the School/Faculty on decolonising and diversifying the curriculum in the future?
Yes
No

Thanks for completing the survey!

### Student survey questions

*Part 1: demographic information*

Q1. Which School is your programme associated with?
Biochemistry
Biomedical Sciences
CMM
PPN

Q2. What year of study are you in?
Foundation year
Year 1
Year 2
Year 3
Year 4

Q3. What is your ethnicity?
Asian/Asian British
Black/African/Caribbean/Black British
White
Mixed/multiple ethnic groups – white and Asian
Mixed/multiple ethnic groups – white and Black
Mixed/multiple ethnic groups – other mixed/multiple ethnic groups, please describe
Not listed – please describe
Prefer not to say

Q4. What is your gender?
Female
Male
Non-binary
Not listed (please describe)
Prefer not to say

Q4a. Is this the gender you were assigned at birth?
Yes
No
Prefer not to say

Q5. What is your sexuality?
Bisexual
Gay/lesbian
Straight/heterosexual
Not listed (please describe)
Prefer not to say

Q6. Do you consider yourself to have a disability?
Yes
No
Prefer not to say

Q7. What is your religion, if you have one?
No religion
Buddhist
Christian
Hindu
Jewish
Muslim

Sikh
Not listed (please describe)
Prefer not to say

*Part 2: decolonising the curriculum*
Q8. How familiar are you with the idea of decolonising the curriculum at university?
Never heard of it
Some awareness
Moderately aware
Very aware
Actively engaged in it

*Decolonisation is broadly about confronting how colonialism, Eurocentrism, and racism have shaped our world. Decolonisation of the curriculum recognises these issues in the science we teach and aims to include a range of perspectives of people who have been overlooked, so as to present a truer picture of our discipline so that it can be more responsive to the needs of all.*

Q9. Are you aware of any activities currently taking place to try to decolonise the curriculum at the following levels?

| | Yes | No |
|---|---|---|
| UK universities | | |
| University of Bristol | | |
| Faculty of Life Sciences | | |
| My School | | |
| My units | | |

Q10. How important do you think it is to decolonise the curriculum in the Biomedical Sciences at the University of Bristol?
Not important 1 2 3 4 5 very important

Q11. To what extent does your curriculum reflect the contribution of diverse cultures to the development of biomedical sciences?
Not at all 1 2 3 4 5 Completely

Q12. To what extent does your curriculum address the needs of diverse populations around the world?
Not at all 1 2 3 4 5 Completely

Q13. To what extent does your curriculum celebrate the contributions of people of colour and other minoritised groups to the development of biomedical sciences?
Not at all 1 2 3 4 5 Completely

Q14. To what extent does your curriculum draw on examples relating to other cultures and people of colour to illustrate ideas/ concepts?
Not at all 1 2 3 4 5 Completely

Q15. What three things would you expect to see in a decolonised curriculum? [Max 280 characters]

*Part 3 – Equality, Diversity and Inclusion in the Biomedical Sciences curriculum*
Q16 Do you feel represented by the science and scientists you learn about in the course?
Not at all 1 2 3 4 5 Completely

Q17. Do you feel that your curriculum is inclusive with respect to the following groups?

| | Yes | No |
|---|---|---|
| People of colour | | |
| People from the Global South | | |
| Women | | |

| |
|---|
| LGBTQIA community |
| Disabled people |
| People of certain religions |
| Other __________________ |

Q18. Do you feel comfortable raising issues about inclusivity, bias or inequality in your course material?

- Very comfortable
- Somewhat comfortable
- Neither comfortable nor uncomfortable
- Somewhat uncomfortable
- Very uncomfortable
- Not aware of any

Q19. To whom would you feel it is best to report these issues if you had them?

- Unit Director
- Individual lecturers
- Personal tutor
- Senior tutor
- Students Union
- Student Reps/SSLC
- University online reporting tool
- Other______________

Q20. If you have experienced any non-inclusive practice (such as bias or inequality) at university, at which level of the organisation was it? Select all that apply:

- Not experienced any non-inclusive practice
- Taught units
- Wider curriculum
- Personal tutoring
- Student societies such as Helix, The Cell, Biomedical Sciences Society, Neuroscience Society etc
- University infrastructure or senior management
- Other______________

Q21 Are there any specific units or topics in which you have experienced non-inclusive practice? Please explain your answer. Please do not name any individuals. If you wish to raise any concerns about unacceptable behaviour, please do so here Report + Support - University of Bristol

Q22. Have you come across any examples of good practice where inequalities have been highlighted and discussed within your course? Please explain them.

Q23 Are there any ways a discussion about equality, diversity and inclusion in science can be encouraged within your course?

*Part 4 - Final comments*
Q24. Do you have any other comments relating to decolonising and diversifying the curriculum?

Q25. Would you be interested in working with the School/Faculty on decolonising and diversifying the curriculum in the future?
Yes
No

Thanks for completing the survey!
